# Categorized analysis of forest ecological values in the China’s conversion cropland to forest program

**DOI:** 10.1101/358960

**Authors:** Wen-Ge Yuan, Jian-Wei Zheng, Jian-Cai Gu, Gui-Qiao Lu

## Abstract

**Background**

The China’s Conversion Cropland to Forest Program (CCFP) is one of the large state ecological construction programs. Up to now, the program has effectively improved the ecological environment and produced large ecological benefit. However, there were also some problems in its implementation process, the program has been sometimes less effective than the expected.

**Methods**

Based on the data and the methods of ‘State report on monitoring ecological effects in CCFP’ and the Chinese Forest Ecosystem Research Network (CFERN) in 2013, we analyzed the categorized ‘forest ecological benefit value’ (B-V) s in the three forest restoration ways in different regions in China to provide references for CCFP construction.

**Results**

The unit area B-Vs in CCFP varied between 35 000 RMBs.hm^−2^.a^−1^ and 100 000 RMBs.hm^−2^.a^−1^. Water conservation B-V and species conservation B-V were the two largest constituents, nutrient accumulation B-V was the least in all the categorized B-Vs on regional and unit area scale. The rank of restoration ways on average unit area total B-Vs was—‘hillside forest conservation’ > ‘returning cropland to forest’ > ‘afforestation on suitable barren hills and wasteland’ in CCFP. Among the categorized B-Vs, some pairs were positively correlated with each other and some were negatively correlative. The correlation coefficients and some regression equations were given in the text and the attached Fig.s.

**Conclusions**

Water conservation B-V was the highest and nutrient accumulation B-V was the lowest whether on regional or unit area scale in CCFP.

Forest ecological B-Vs varied in different forest restoration ways and different regions in CCFP. The ‘hillside forest conservation’ restoration way and the water conservation B-V should be paid more attention in China’s future forest restoration. We suggest that suitable forest restoration ways should be selective according to the regional specific and ecological targets.

There were correlations among the categorized B-Vs, and the correlations varied with different forest restoration ways in CCFP. Knowing about the correlations could clarify the targeted restoration ways according to the goal of ecological benefit.

## 1. Background

Since the 20^th^ century, a series of ecological problems have become more serious (Zhou 2008) with the increasing population (Wikipedia 2004), irrational development and utilization of natural resources, such as deforestation, biodiversity loss, soil erosion, desertification and so on (Pandit et al. 2007; Sims et al. 1996). The ecological problems have attracted worldwide attention (Wu et al. 2009). Ecosystem service function becomes the hotspot and frontier of current international research (Bailey 1998, 2014; Beier et al. 2010; Zhang et al. 2010; Higgins et al. 2005). As a critical component of ecosystem, forest ecosystem plays important roles in water and soil conservation (So-co), carbon fixation and oxygen release (Cf-Or), nutrient accumulation (Nu-ac), atmosphere purification (At-pu), biological diversity protection and so on (Wang et al. 2009; Metzger et al. 2005). To discuss the service functions of forest ecosystem, a lot of researches (Jonge et al. 2012; Li et al. 2009; Zhang et al. 2004, 2001; Liu et al. 2003, 1996; Zhang et al. 1988; Zhou et al. 1995) and practices have been carried out and many valuable results have been applied to improve the environmental quality. Some scientists divided the forests into northern and southern types to study the forest service functions (Constanza et al. 1997). Capotorti et al. (2012) discussed the ecological classification of land and conservation of biodiversity in Italy. Niu et al. (2012) studied categorized forest ecological values and concluded that the percentages of water and soil values were 40.51% and 9.91%, respectively, in Chinese forest ecosystem.

In order to improve the situation of serious ecological deterioration, the largest ecological program—CCFP has been implemented in China since 1999. It is of great importance in improving the state ecological environment, preventing water and soil erosion, improving water conservation (Wa-co) ability, speeding up the adjustment of rural industrial structure, increasing the overall agricultural capacity, promoting the harmonious development between human beings and nature. Although the program has made great achievement, there were also many problems in the implementation (Yang et al. 2011). For example, the program has been sometimes less effective than the expected. (Cao et al. 2011). Understanding characteristics of categorized B-Vs in different forest restoration ways and different regions could make the afforestation more efficient for the future implementation in CCFP.

For the purpose of more systematical observation and study on the functions of the forest ecosystem, CFERN has been established in China since the end of 1950s (Wang et al. 2010). By 2015, the number of ecological observation stations had reached 110 in CFERN. Nearly 100 ecological indicators including atmosphere, soil, forest and creatures were involved in the range of the observation. This would strongly promote the state ecological construction. Whereas, categorized comparative analyses and correlation studies for B-Vs in different restoration ways on a large scale and in long-term in CCFP were rarely reported.

Based on the data of ‘State report on monitoring ecological effects in CCFP’ and CFERN in 2013, we carried on the categorized analysis on B-Vs in CCFP. The objective of the study is to find the differences, features and the relationships of the categorized B-Vs among the three forest restoration ways—‘hillside forest conservation’ (H-f-c), ‘afforestation on suitable barren hills and wasteland’ (A-b-w), ‘returning cropland to forest’ (R-c-f). We hope it will be able to provide references for the construction of forest restoration and be an interesting issue for us to communicate with the peers.

## 2. Methods

### 2.1. Classifications of B-Vs and restoration ways

We divided the B-Vs into six categories: Water conservation; Soil conservation; Carbon fixation and oxygen release; Nutrient accumulation; Atmosphere purification and species conservation. The forest restoration ways were classified as three kinds—‘hillside forest conservation’, ‘returning cropland to forest’, ‘afforestation on suitable barren hills and wasteland’.

### 2.2. Original data and study regions

The data of ‘State report on monitoring ecological effects in CCFP in 2013’ (China’s State Forestry Administration 2013) and the Chinese Forest Ecosystem Research Network (CFERN) in 2013 were cited in this study. All the observation stations, which the evaluation data came from, are under the technical standards and management regulations of the observation and evaluation of ecological effects in CCFP (CFERN 2013). Six key ecological monitoring provinces and the relevant zones, which represent the main biotope types in CCFP, were involved in the analysis. They are Hebei province (HE-B), Liaoning province (L-N), Hubei province (HU-B), Hunan province (HU-N), Yunnan province (Y-N) and Gansu province (G-S). The representative zones are: Shijiazhuang city, Tangshan city and Qinhuangdao city in HE-B; Shenyang city and Panjin city in L-N; Shiyan city, Xiaogan city and Enshi Tujia and Miao autonomous prefecture in HU-B; Huaihua city, Hengyang city and Xiangxi Tujia and Miao autonomous prefecture in HU-N; Kunming city and Dali city in Y-N; Lanzhou city and Gannan Tibetan autonomous prefecture in G-S.

### 2.3. Discounted ecological value

Parameters of ecological value were discounted to 2013. They are calculated as follows:

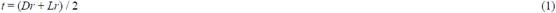

*t*: deposit and loan equilibrium interest rate

*Dr*: average deposit rate

*Lr*: average loan rate

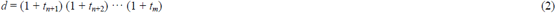

*d*: discount rate

*t*: deposit and loan equilibrium interest rate

*n*: the year of obtained parameters

*m*: the year of evaluation

### 2.4. Data processing

Observation Methodology for Long-term Forest Ecosystem Research (*LY/T1952–2011*) (China’s State Forestry Administration 2011), Indicators System for Long-term Observation of Forest Ecosystem (*LY/T1606–2003*) (China’s State Forestry Administration 2003), Specifications for Assessment of Forest Ecosystem Services in China (*LY/T1721–*2008) (China’s State Forestry Administration 2008) and the calculation formulas in ‘State report on monitoring ecological effects in CCFP in 2013’ (China’s State Forestry Administration 2013) were exploited in the B-V calculated process. The gained B-Vs and SPSS 19.0 were used in the analysis.

## 3. Nature conditions of study regions

The locations of the six key ecological monitoring provinces are shown in Fig. 1.

**Fig. 1.**
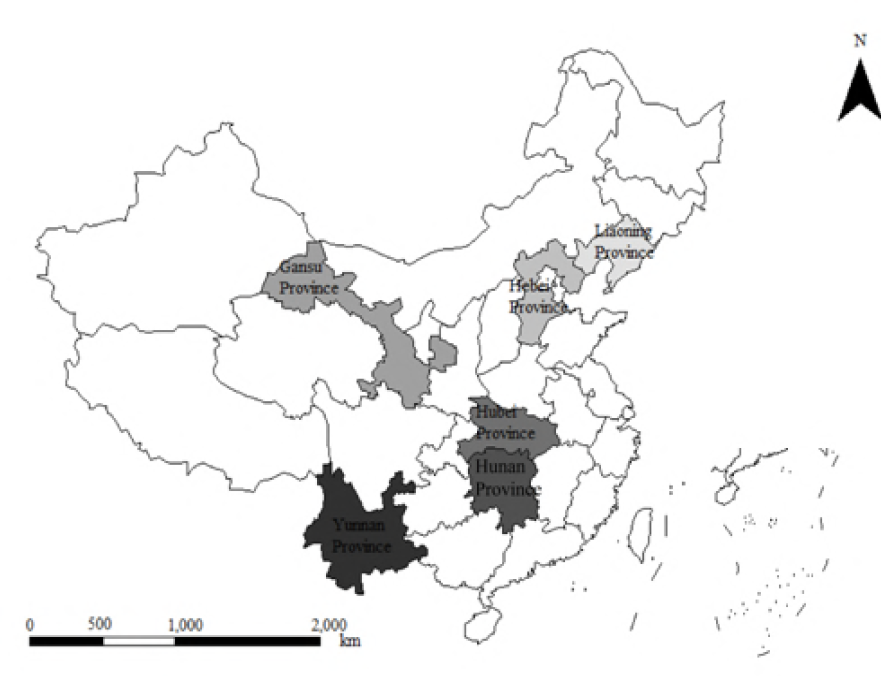
The six key monitoring provinces, China.

HE-B is located in 113°27′ E–119°50′ E, 36°05′ N–42°40′ N with total area of 188 500 km^2^. The terrain is downward from northwest to southeast. The northwest part is mainly mountainous and hilly land, the central and southern parts are plain. The average annual precipitation is 484.5 mm and the average annual temperature is −0.5°C–14.2°C (China Meteorological Administration 1981–2010). Soil types are predominantly red and yellow earths. Forests consist of coniferous, broad-leaved, mixed forest and shrub.

L-N is located in 118°53′ E–125°46′ E, 38°43′ N–43°26′ N with total area of 148 000 km^2^. The terrain is downward from the north to the south. The east and west parts are mainly mountain and hilly lands, the central part is plain. The average annual precipitation is 660.0 mm and the average annual temperature is 8.3°C (China Meteorological Administration 1981–2010). Soil types are mainly dark-brown earths and solonetzs. Forest types are mainly coniferous, broad-leaved, mixed forest and shrub.

HU-N is located in 108°48′ E–114°15′ E, 30°08′ N–24°38′ N, mainly consists of low mountain and hills with total area of 211 800 km^2^. The terrain is high in the south and low, flat in the center and north. The average annual precipitation is 1 200.0 mm–1 700.0 mm and the average annual temperature is 15°C–18°C (China Meteorological Administration 1981–2010). Soil types are mainly red or yellow earths. Forest types are mainly coniferous, broad-leaved, mixed and bamboo forest. HU-B is located in 108°21′ E–116°07′ E, 29°05′ N–33°20′ N, consists of northern mountain land, hills and plain in the center and south with total area of 185 900 km^2^. The average annual precipitation is 800.0 mm–1 600.0 mm and the average annual temperature is 15°C–17°C (China Meteorological Administration 1981–2010). Soil types are mainly red or yellow earths. Forest types are mainly coniferous, broad-leaved, mixed and bamboo forest.

Y-N is located in 97°52′ E–106°18′ E, 21°13′ N–29°25′ N, consists of mountain, hills, basin and plateau with total area of 390 000 km^2^. The average annual precipitation is 1 100.0 mm–1 600.0 mm and the average annual temperature is 5°C–24°C (China Meteorological Administration 1981–2010). Soil types are mainly red or yellow earths with part of Grey-cinnamon soils. Forest types are mainly coniferous, broad-leaved, mixed and bamboo forest.

G-S is located in 92°13′ E–108°46′ E, 32°31′ N–42°57′ N, consists of staggered mountain, valley and plain with total area of 425 900 km^2^. The average annual precipitation is 386.0 mm and the average annual temperature is 0°C–15°C (China Meteorological Administration 1981–2010). Soil type is mainly Aeolian sandy soil. Forest types are mainly coniferous, broad-leaved, mixed and shrub.

## 4. Results

### 4.1. Area distribution of different forest restoration ways

From 1999 to 2013, the total forest restoration area in CCFP reached 29 819 100 hm^2^. ‘Afforestation on suitable barren hills and wasteland’ accounted for 17 455 000 hm^2^, ‘returning cropland to forest’ accounted for 9 264 133 hm^2^ and ‘hillside forest conservation’ accounted for 3 100 000 hm2. HE-B and G-S formed larger forest restoration areas of 1 866 700 hm^2^ and 1 896 900 hm^2^, respectively (Fig.2).

**Fig. 2.**
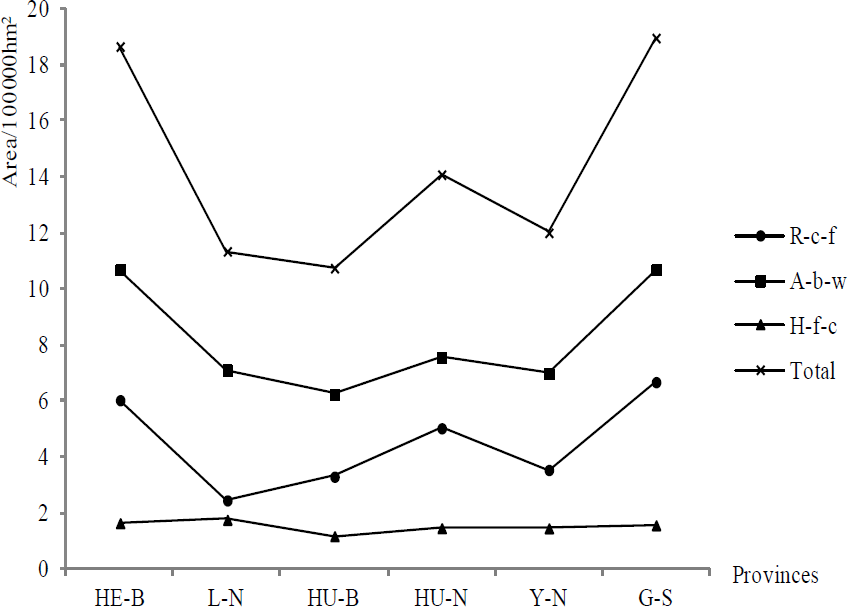
Forest areas of different vegetation restoration ways in key ecological monitoring provinces in CCFP.

### 4.2. B-Vs of ‘hillside forest conservation’

### 4.2.1. B-V features of ‘hillside forest conservation’

‘Hillside forest conservation’ is a forest restoration way to avoid human destruction and facilitate natural reforestation through regular hillside-closing measures in suitable mountain regions. Totally 3 100 000 hm^2^ of ‘hillside forest conservation’ in CCFP had been formed until 2013.

For this way of conservation, it was shown that the water conservation B-V was higher than the other categorized B-Vs whether on regional or on unit area scale in the ecological monitoring provinces. The water conservation B-V accounted for approximately 46.6% (HE-B had the highest percentage of 57.6%, L-N had the lowest percentage of 31.7%) of the total B-V on regional scale and approximately 46.5% (annual average, 28 287.42 RMBs·hm^−2^.a^−1^ / 60 814.01 RMBs·hm^−2^.a^−1^) on unit area scale. The water conservation B-V was obviously higher in HE-B with unit area B-V of 55 076.01 RMBs·hm^−2^.a^−1^. The species conservation B-V ranked the second except in G-S where produced the highest soil conservation B-V with unit area B-V of 8 205.44 RMBs·hm^−2^.a^−1^, and it was higher in the southern provinces (e.g. HU-N, HU-B and Y-N) whether in regions or in unit areas. For example, HU-N produced the highest species conservation B-V with unit area B-V of 25 150.16 RMBs·hm^−2^.a^−1^, and the northwest province—G-S produced only 3713.04 RMBs·hm^−2^.a^−1^. The nutrient accumulation B-V was the lowest in all the categorized B-Vs with an average of 928.85 RMBs·hm^−2^.a^−1^. The lowest ‘carbon fixation and oxygen release’ B-V occurred in HU-N with unit area B-V of 670.94 RMBs·hm^−2^.a^−1^.

There were also different performances between regional and unit area scale. The rank of regional total B-V was HE-B > Y-N > HU-N > HU-B > L-N > G-S, whereas, the rank of unit area total B-V was HE-B > HU-B > Y-N > HU-N > L-N > G-S. L-N had formed the highest regional annual soil conservation B-V with 1 322.00 million RMBs, but its unit area soil conservation B-V ranked the second with 7 457.01 RMBs·hm^−2^.a^−1^. HU-N ranked the second regional water conservation B-V with 4322.00 million RMBs, meanwhile, its unit area water conservation B-V ranked the third with 29216.48 RMBs·hm^−2^.a^−1^ (Fig.3).

**Fig. 3.**
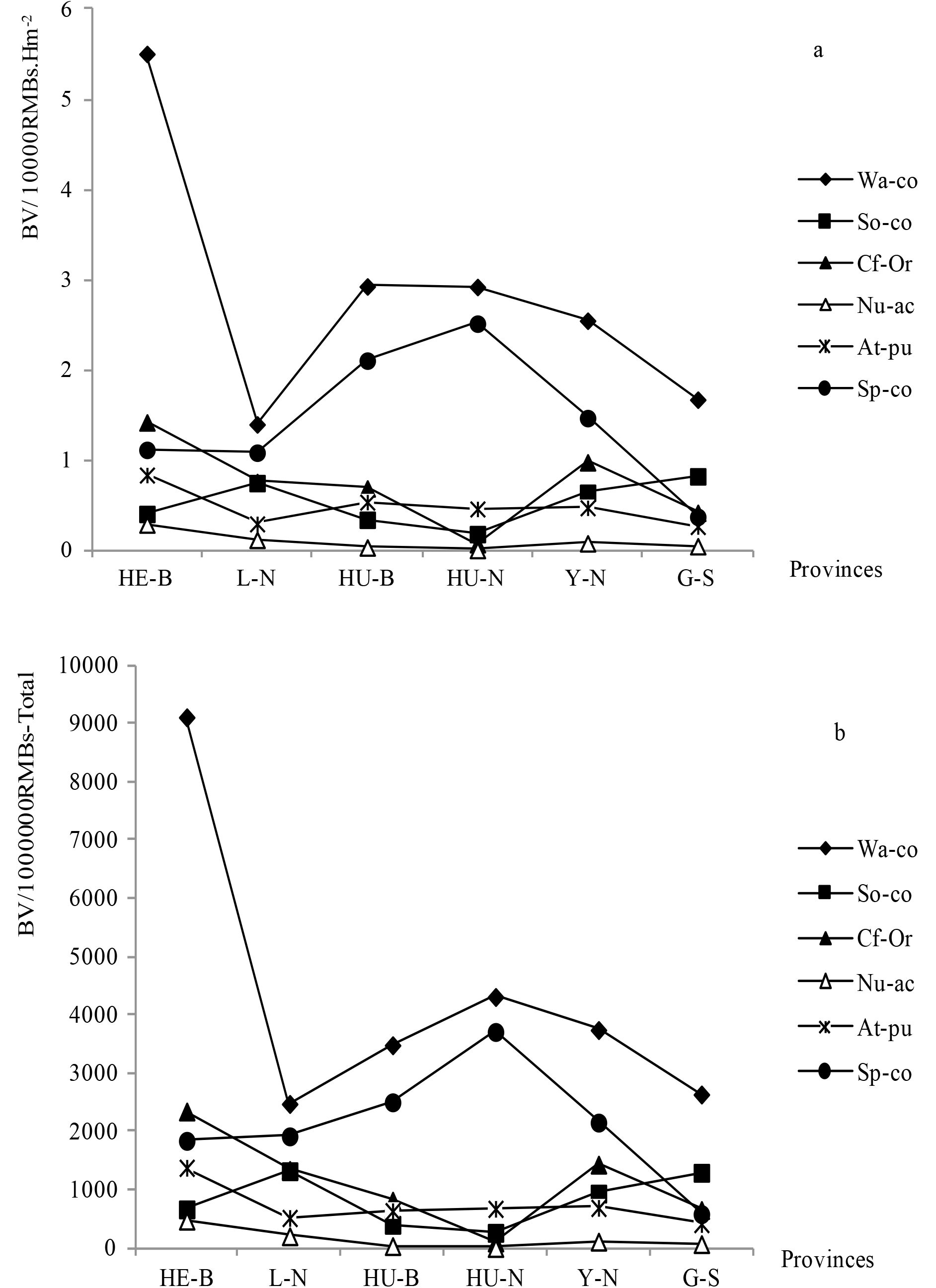
Annual B-Vs of hillside forest conservation in different provinces a: For unit area; b: For regional area.

### 4.2.2. Correlation analysis for ‘hillside forest conservation’ B-Vs

To find the relationships among the ‘hillside forest conservation’ B-Vs, fourteen data lines of categorized unit area B-Vs that come from the six monitoring provinces in 2013 were calculated. The result showed that the unit area water conservation B-V and the unit area atmosphere purification B-V had significantly positive correlations with their total B-Vs (r=0.906, p<0.01; r=0.914, p<0.01), so were the water conservation B-V with atmosphere purification B-V (r=0.722, p<0.01) and water conservation B-V with nutrient accumulation B-V (r=0.633, p<0.01). Meanwhile, the unit area nutrient accumulation B-V had significantly negative correlation with the unit area species conservation B-V (r=-0.532, p<0.05). We adopted 77 data lines of categorized regional B-Vs to analyze their correlations. The result showed that regional total B-V had significantly positive correlations with all the relevant categorized regional B-Vs (Table 1). Several regressions among categorized B-Vs of ‘hillside forest conservation’ were shown in Fig. 4.

**Table 1.**
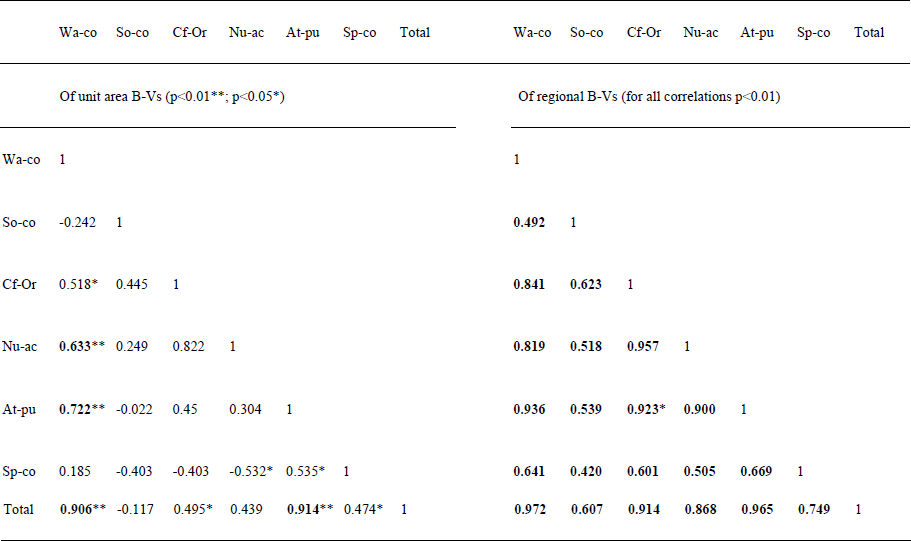
Correlation coefficients among the annual B-Vs of hillside forest conservation.

**Fig. 4.**
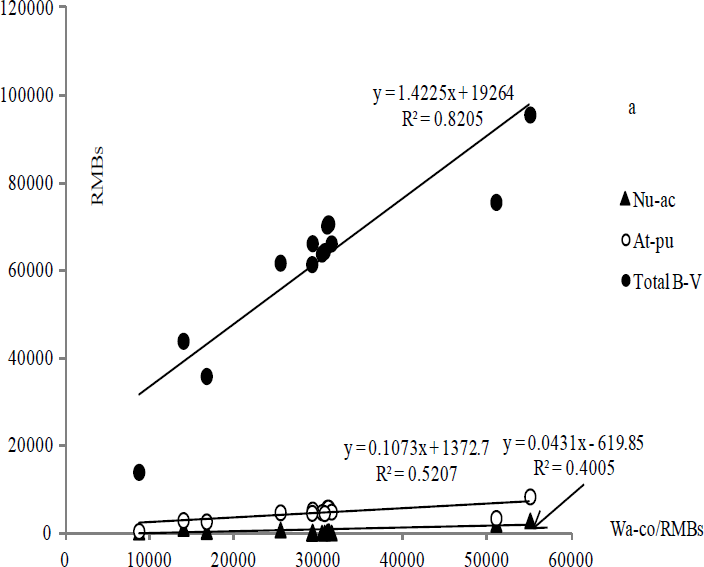
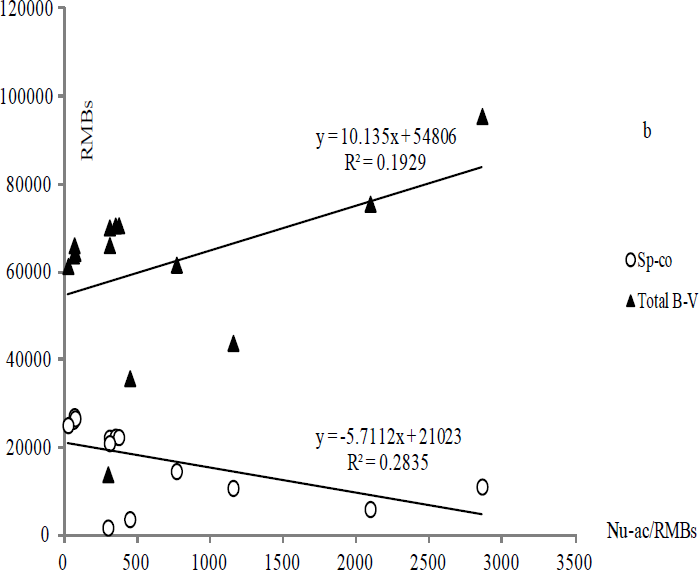
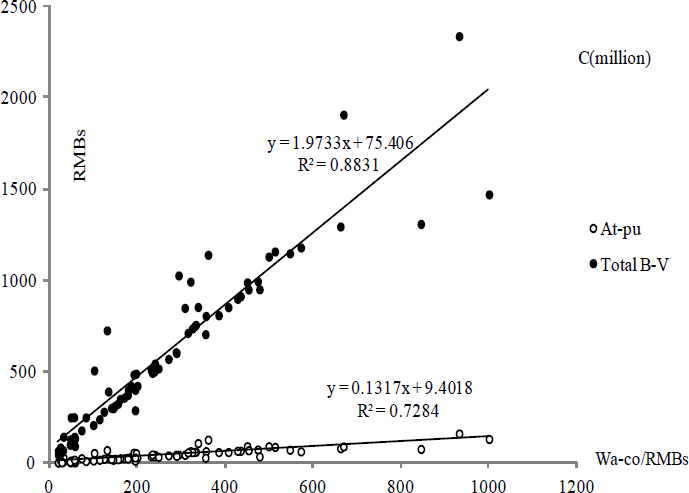
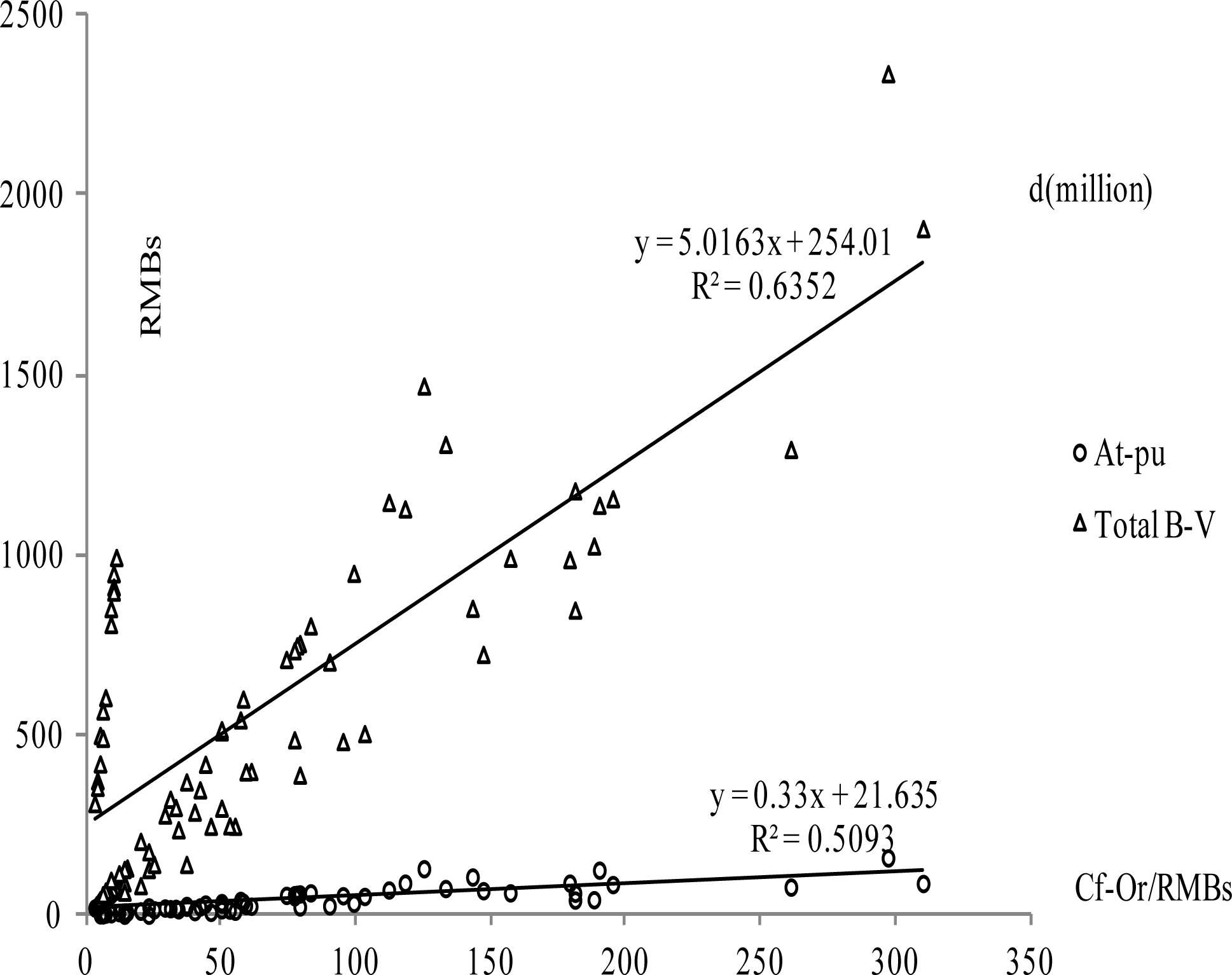
The relationships among annual B-Vs of hillside forest conservation a, b: For unit areas; c, d: For regional areas.

### 4.3. B-Vs of ‘returning cropland to forest’

#### 4.3.1. B-V Features of ‘returning cropland to forest’

According to the state regulation (China’s State Forestry Administration 2002), the ‘returning cropland to forest’ was performed in the regions with serious soil and water loss, desertification and stony desertification, salinization; The low yield regions with crucial ecological function; The regions at river source or river side and the croplands with ecological importance which were seriously damaged by the wind and sand.

By 2013, China had finished 9 264 133 hm^2^ of ‘returning cropland to forest’. It was shown that water conservation was the main part in the relevant total B-V, which accounted for approximately 49.0% (HE-B had the highest percentage of 58.3%, L-N had the lowest percentage of 30.0%) of the total B-V on regional scale and approximately 47.3% (annual average, 25 238.93 RMBs·hm^−2^.a^−1^ / 53 386.17 RMBs·hm^−2^.a^−1^) on unit area scale. The nutrient accumulation B-V was still the lowest with the unit area B-V of 781.06 RMBs·hm^−2^.a^−1^. Obviously, HU-N had the highest species conservation B-V and the lowest ‘carbon fixation and oxygen release’ B-V whether on regional scale or on unit area scale. G-S produced the highest soil conservation B-V with unit area B-V of 9 305.57 RMBs·hm^−2^.a^−1^.

The rank of regional total B-V was HU-N > HE-B > G-S > Y-N > HU-B > L-N, and the rank of unit area total B-V was HU-N > Y-N > HE-B > HU-B > G-S > L-N. HE-B produced the highest regional annual water conservation B-V with 19 228.00 million RMBs, but its unit area water conservation B-V ranked the second with 31 811.40 RMBs·hm^−2^.a^−1^. In addition, HE-B produced the highest regional annual B-V of ‘carbon fixation and oxygen release’ with 5 410.00 million RMBs, and its unit area B-V of ‘carbon fixation and oxygen release’ ranked the third with 8 950.47 RMBs·hm^−2^.a^−1^ (Fig. 5).

**Fig. 5.**
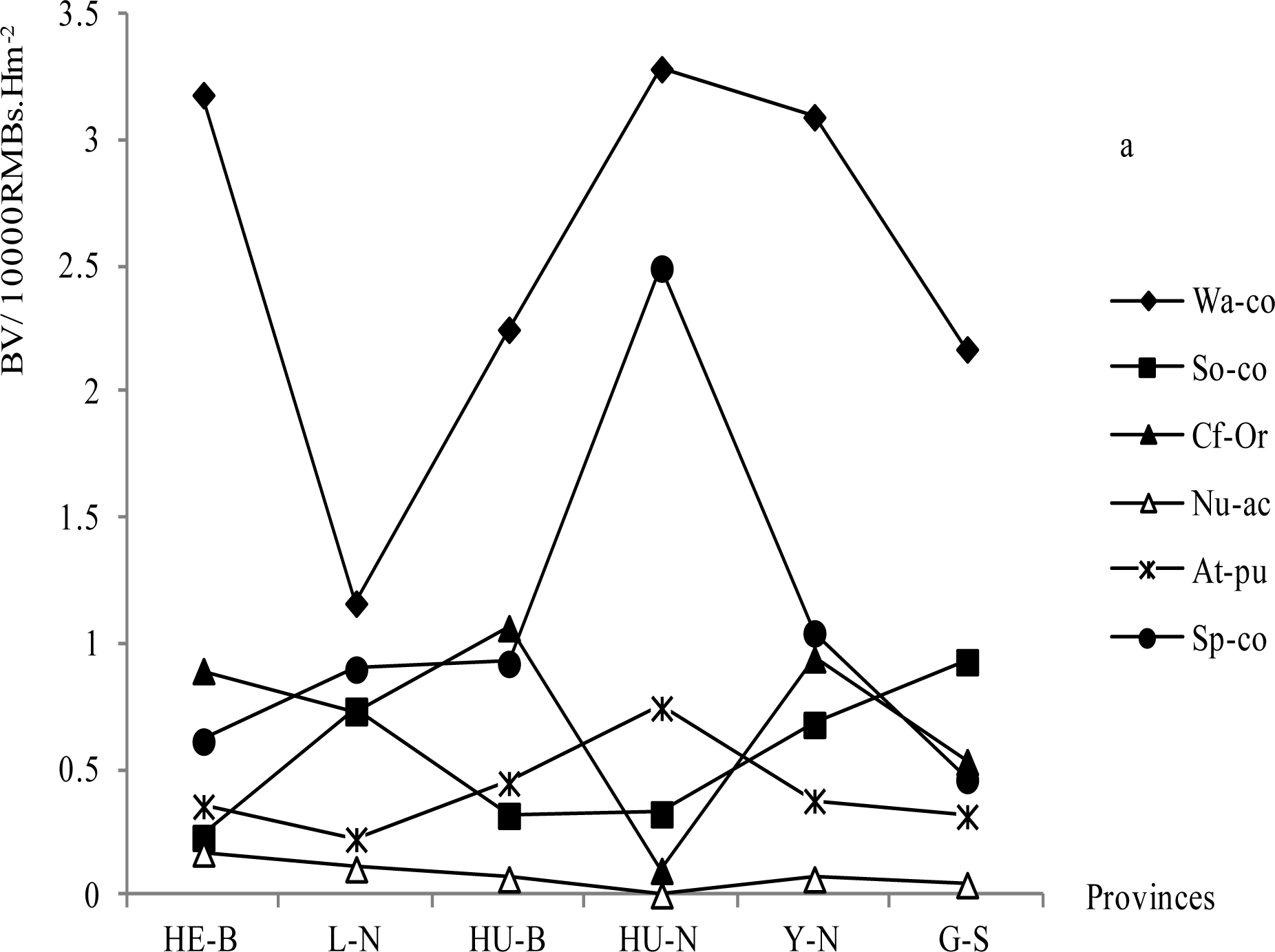
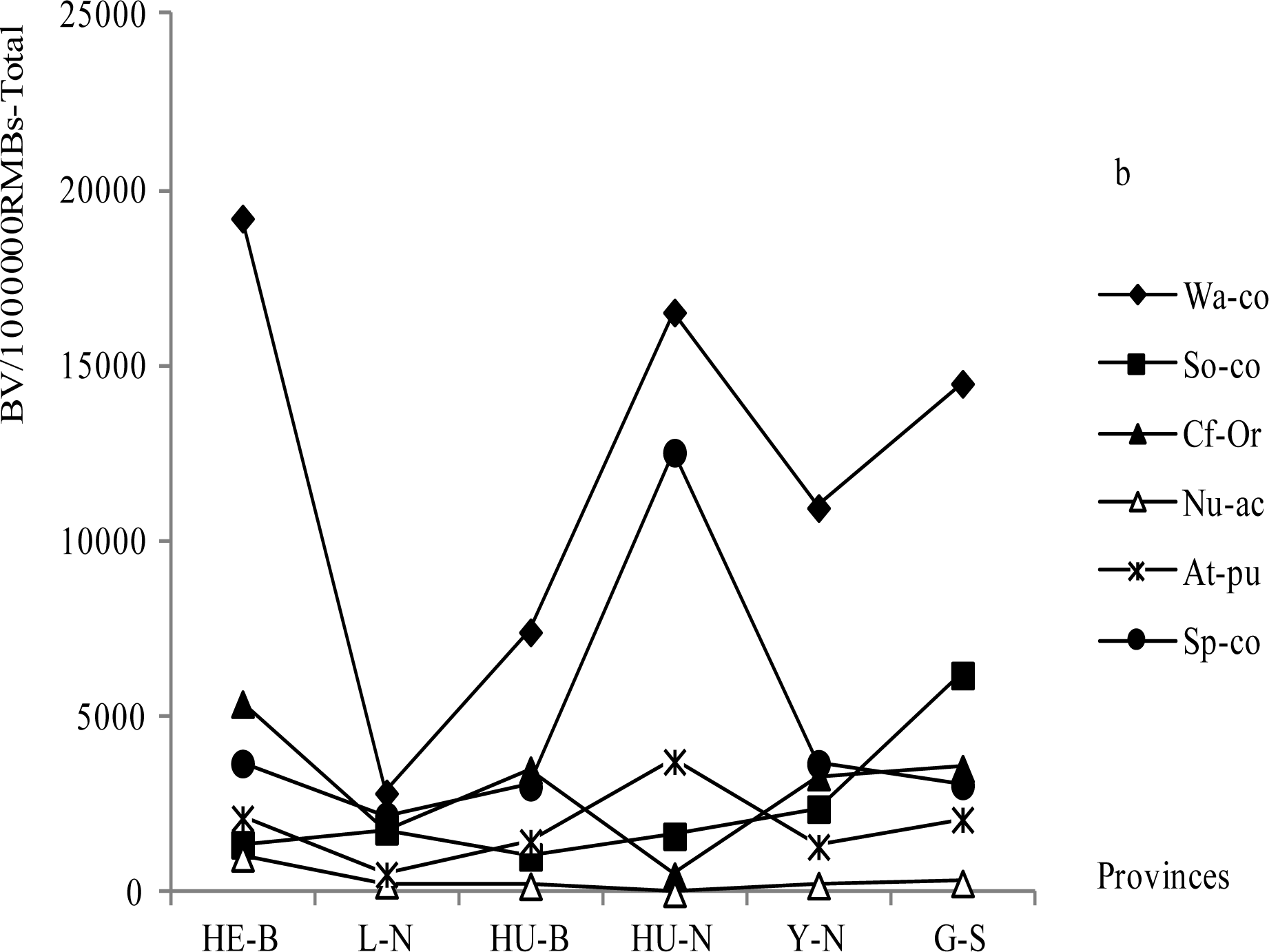
Annual B-VS in the returning cropland to forest way in different provinces a: For unit areas; b: For regional areas.

#### 4.3.2. Correlation analysis for B-Vs of ‘returning cropland to forest’

Thirty-one data lines of categorized regional B-Vs and fourteen data lines of categorized unit area B-Vs in the ‘returning cropland to forest’ way were analyzed. The result showed that some of the correlations among the categorized B-Vs were positive and others were negative whether on regional scale or on unit area scale. However, the consistent result showed that the B-V pairs of water conservation with its total, ‘carbon fixation and oxygen release’ B-V with its relevant nutrient accumulation B-V, atmosphere purification B-V with its relevant species conservation B-V, atmosphere purification B-V with its total, and species conservation B-V with its total had significantly positive correlations. Conversely, the B-V pair of nutrient accumulation B-V with its relevant species conservation B-V was negatively correlative (Table 2). Several regressions of the B-Vs were shown in Fig. 6.

**Fig. 6.**
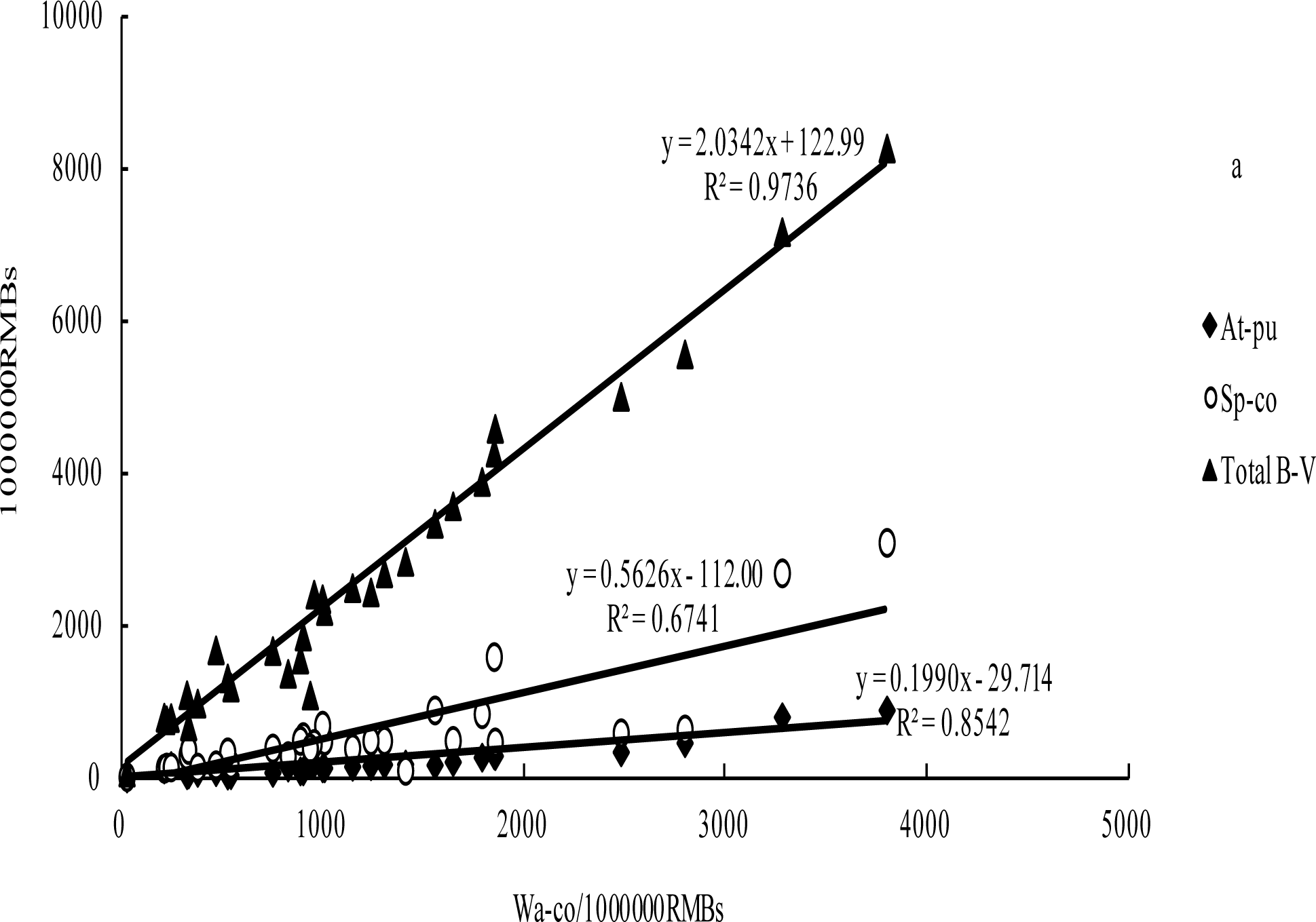
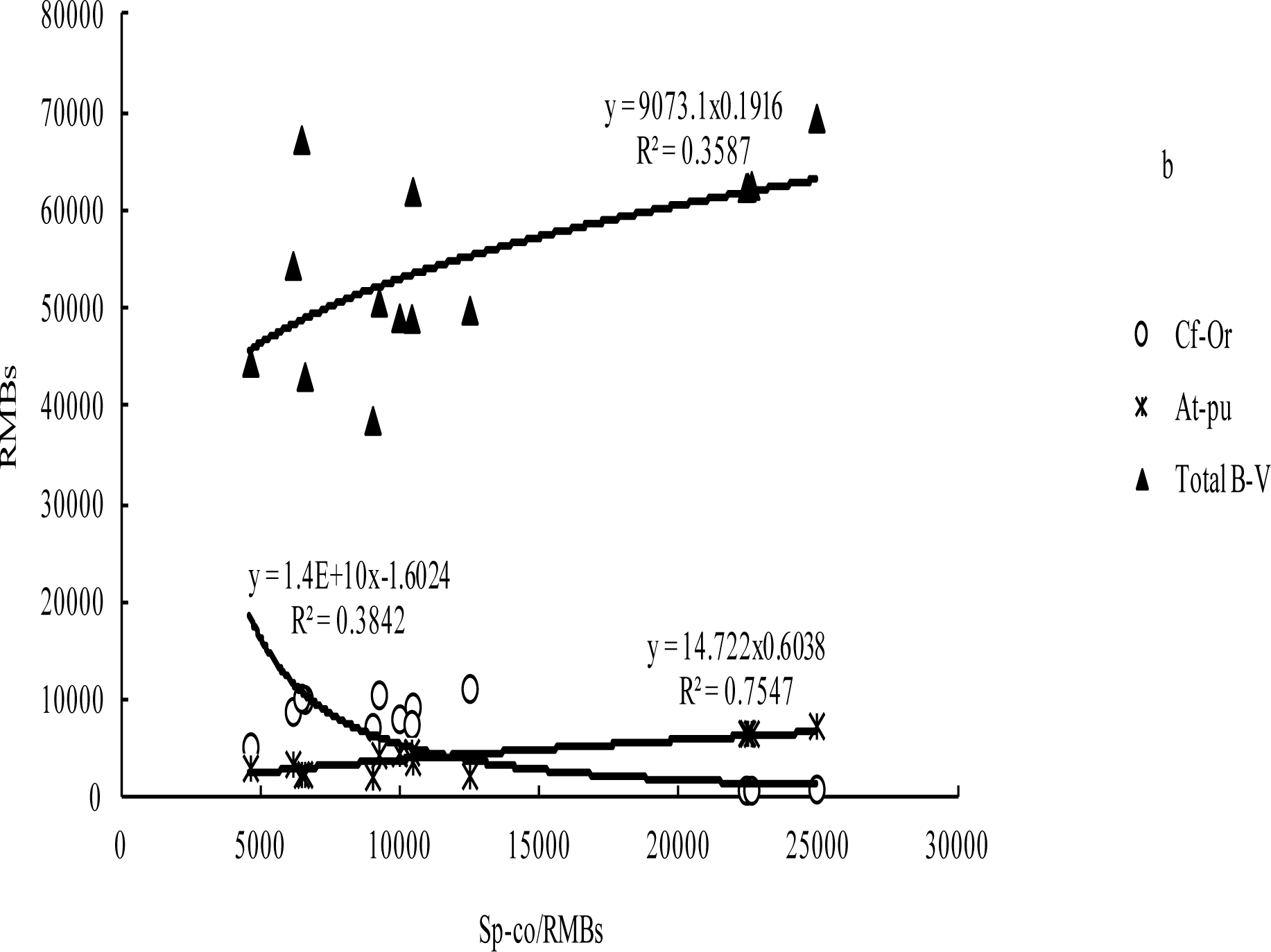
The relationships among annual B-Vs of returning cropland to forest a: For regional areas; b: For unit areas.

**Table 2.**
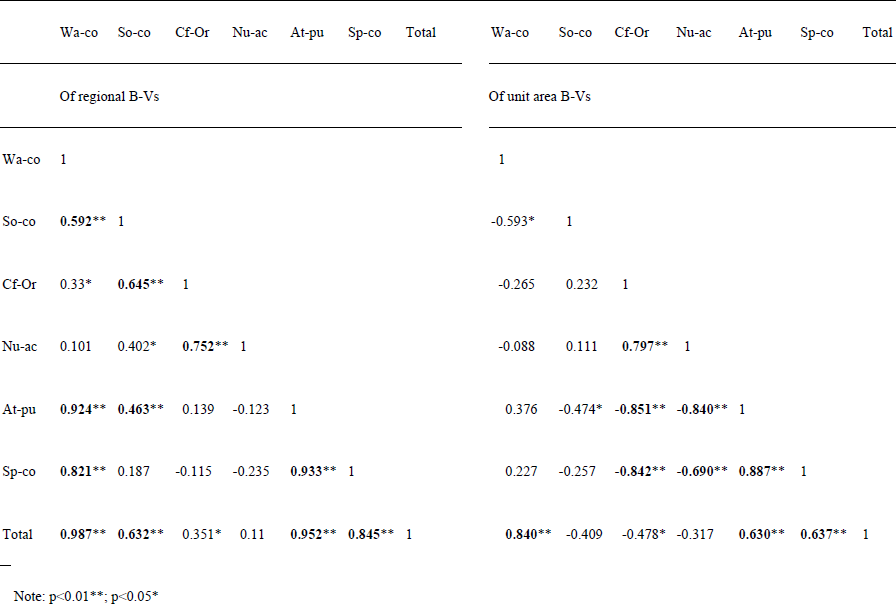
Correlation coefficients among the annual B-Vs of returning cropland to forest.

### 4.4. B-Vs of ‘afforestation on suitable barren hills and wasteland’

#### 4.4.1. B-V Features of ‘afforestation on suitable barren hills and wasteland’

By 2013, the total area of ‘afforestation on suitable barren hills and wasteland’ had reached 17 455 000 hm^2^. In the way of ‘afforestation on suitable barren hills and wasteland’, the categorized B-Vs had the similar ranking trends between regional scale and unit area scale among the monitoring provinces. Water conservation B-V was still the largest in all the B-Vs, which accounted for 45.7% (HE-B had the highest percentage of 58.5%, L-N had the lowest percentage of 24.7%) of the total B-V on regional scale and approximately 44.9% (annual average, 22 950.56 RMBs·hm^−2^.a^−1^ / 51 082.92 RMBs·hm^−2^.a^−1^) on unit area scale. Nutrient accumulation B-V was the lowest with the unit area B-V of 4 877.82 RMBs·hm^−2^.a^−1^. Atmosphere purification B-Vs varied little among the monitoring provinces. G-S produced the highest soil conservation B-V with regional annual 12 414.00 million RMBs, and unit area of 11 599.37 RMBs·hm^−2^.a^−1^. HU-N possessed the highest species conservation B-V and the lowest ‘carbon fixation and oxygen release’ B-V (Fig. 7).

**Fig. 7.**
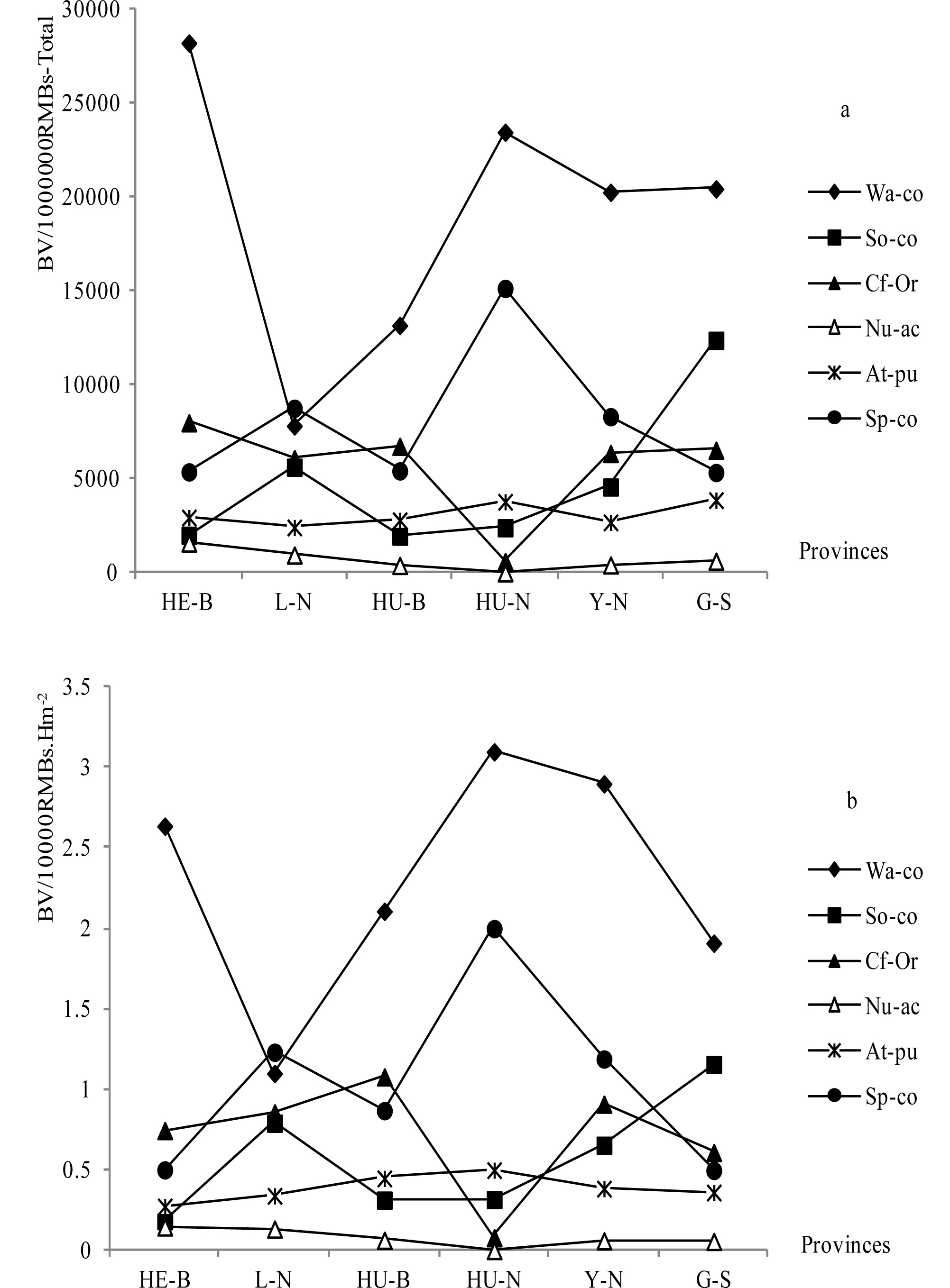
Annual B-Vs of afforestation on suitable barren hills and wasteland in different provinces a: For regional area; b: For unit area.

The rank of regional total B-V was G-S > HE-B > HU-N> Y-N > L-N > HU-B, meanwhile, the rank of unit area total B-V was Y-N > HU-N > HU-B > G-S > HE-B > L-N. HU-N produced the highest unit area water conservation B-V of 31 006.82 RMBs·hm^−2^.a^−1^, but its regional annual ranked the second with 23492.00 million RMBs. HU-B formed the highest unit area of ‘carbon fixation and oxygen release’ B-V with 10 826.46 RMBs·hm^−2^.a^−1^, and its regional annual ranked the second with 6 771.00 million RMBs.

#### 4.4.2. Correlation analysis for B-Vs of ‘afforestation on suitable barren hills and wasteland’

We analyzed 14 data lines of unit area B-Vs and 31 data lines of regional B-Vs of ‘afforestation on suitable barren hills and wasteland’. For regional B-Vs, The results suggested that every categorized regional B-V had significantly positive correlations with other relevant B-Vs except the four pairs: species conservation with soil conservation, species conservation with ‘carbon fixation and oxygen release’, species conservation with nutrient accumulation and atmosphere purification with nutrient accumulation. For unit area B-Vs, the following pairs had significantly positive correlations at p<0.01 levels: water conservation with its total, ‘carbon fixation and oxygen release’ with its relevant nutrient accumulation, atmosphere purification with its relevant species conservation, and species conservation with its total. On the contrary, some pairs had significantly negative correlations at p<0.01 levels, including ‘carbon fixation and oxygen release’ with its relevant species conservation, ‘carbon fixation and oxygen release’ with its relevant atmosphere purification, nutrient accumulation with its relevant species conservation, and nutrient accumulation with its relevant atmosphere purification (Table 3). Several regressions were given in Fig. 8.

**Fig. 8.**
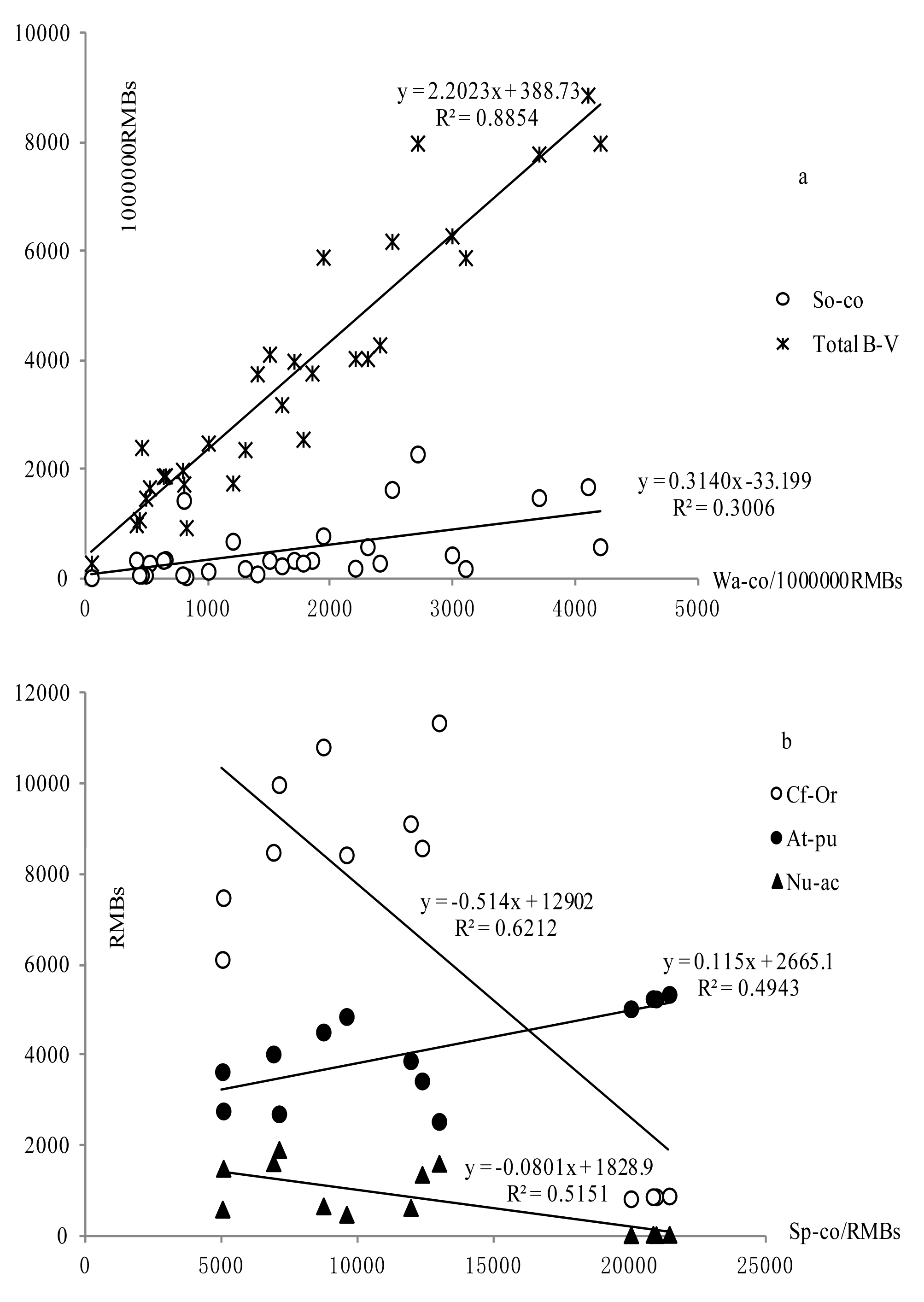
The relationships among B-Vs of afforestation on suitable barren hills and wasteland a: For regional area; b: For unit area.

**Table 3.**
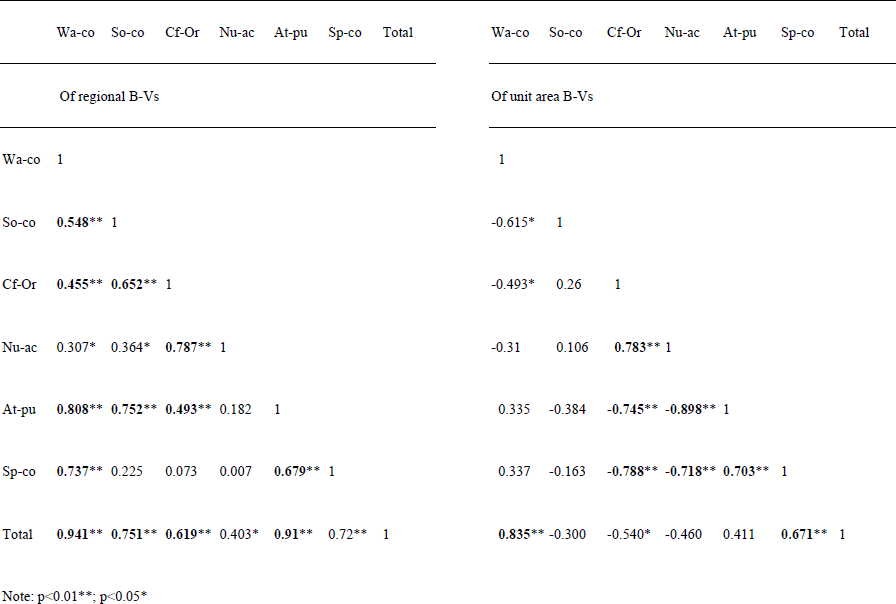
Correlation coefficients among the annual B-Vs of afforestation on suitable barren hills and wasteland.

### 4.5. Comparisons of annual categorized unit area B-Vs in different regions and forest restoration ways

Categorized unit area B-Vs were calculated in different forest restoration ways (Fig. 9). For the water conservation B-Vs, there were no obviously differences among the three restoration ways in the monitoring provinces expect HE-B, in which the water conservation B-V of ‘hillside forest conservation’ was obviously higher than the B-Vs of other two restoration ways. The soil conservation B-Vs of G-S, L-N and Y-N were higher than other provinces in all the three restoration ways. For the ‘carbon fixation and oxygen release’ B-V, the ‘hillside forest conservation’ was higher than the other two ways in HE-B, and there were no obviously differences among the three ways in other provinces. The ‘carbon fixation and oxygen release’ B-Vs of the three ways in HU-N and G-S were lower than other provinces especially in HU-N. HE-B produced higher nutrient accumulation B-V in the ‘hillside forest conservation’ way. L-N and G-S had higher nutrient accumulation B-Vs in the way of ‘afforestation on suitable barren hills and wasteland’. HU-N had the least nutrient accumulation B-V, but its species conservation and the atmosphere purification B-Vs were obviously higher in all the three restoration ways. For species conservation B-V, HU-B, HE-B and Y-N produced higher B-Vs in ‘hillside forest conservation’ way. HE-B produced higher ‘hillside forest conservation’ B-Vs. HU-N produced higher atmosphere purification B-V in ‘returning cropland to forest’ way. Except soil conservation B-V, ‘hillside forest conservation’ had higher categorized B-Vs especially in water conservation B-V, atmosphere purification B-V and species conservation B-V. ‘Afforestation on suitable barren hills and wasteland’ produced more soil conservation B-V (Table 4).

**Fig. 9.**
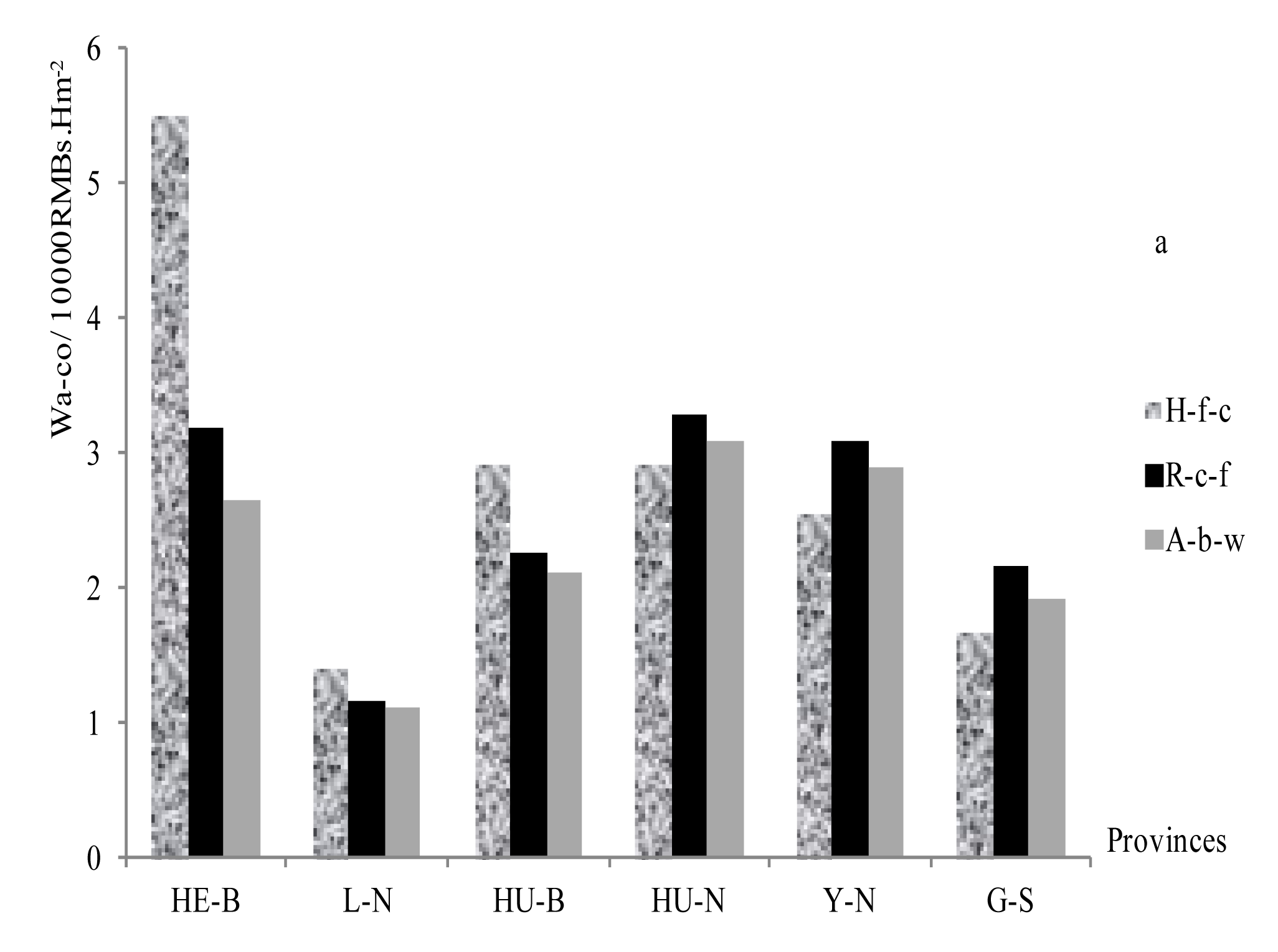
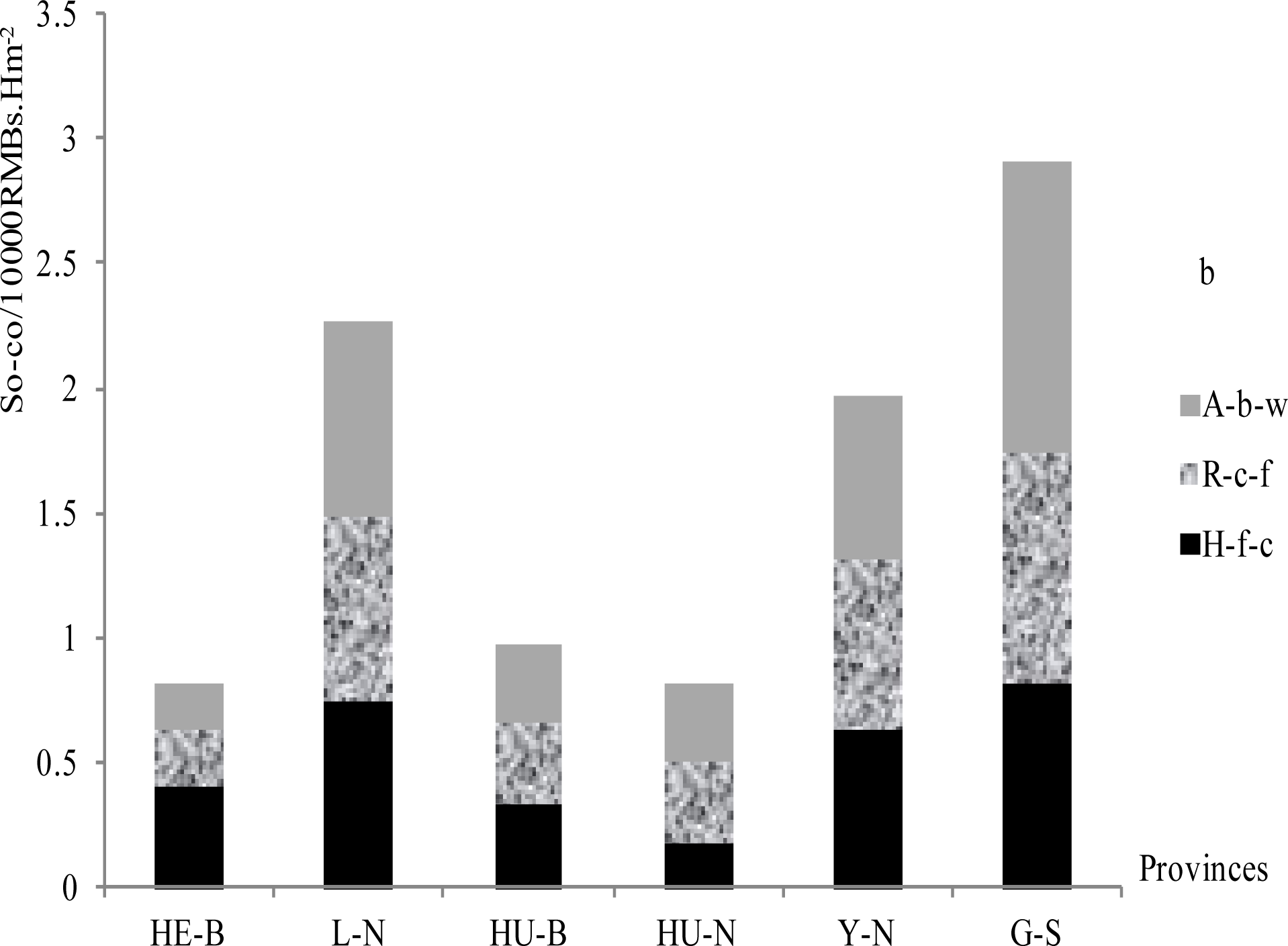
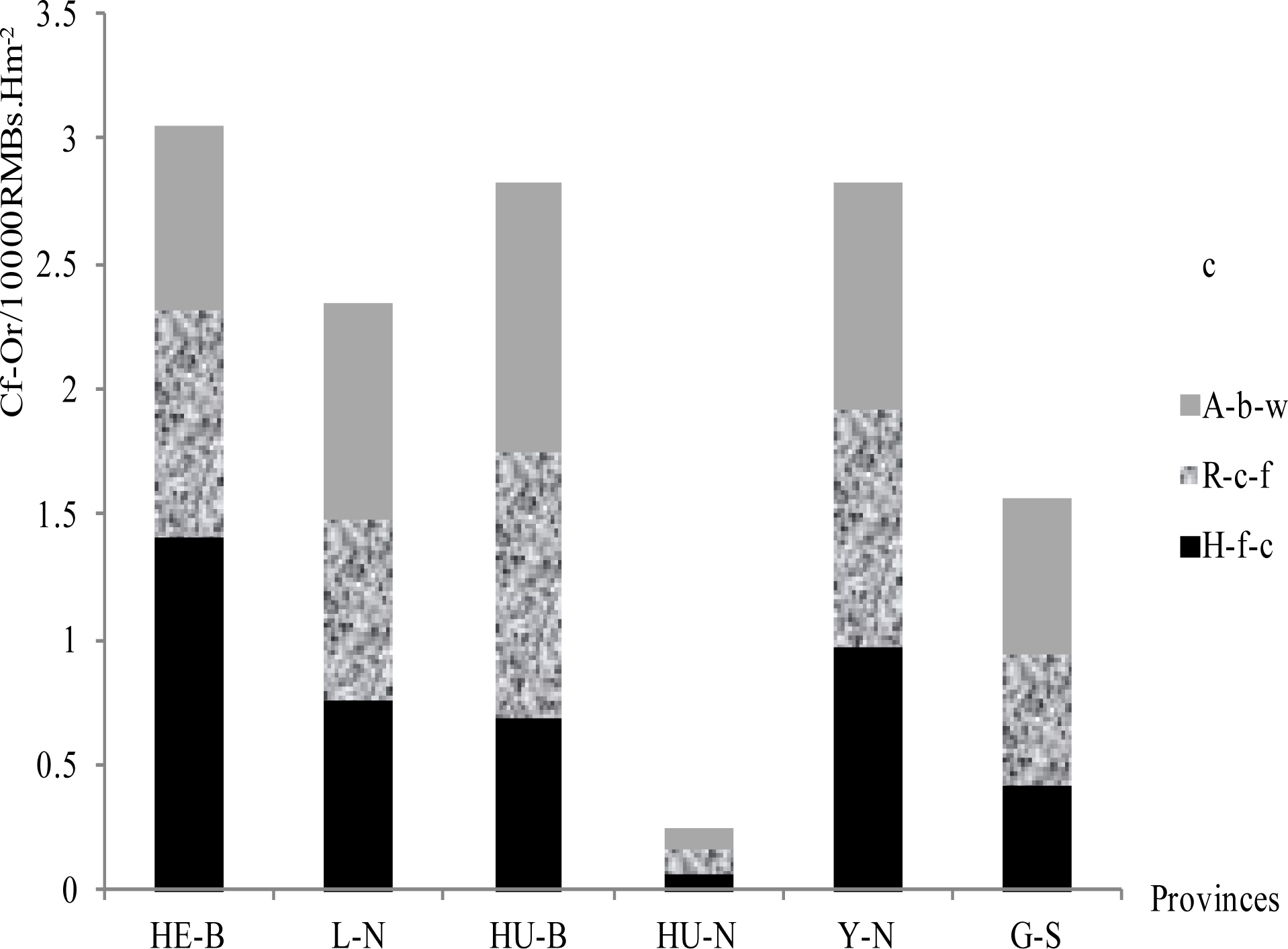
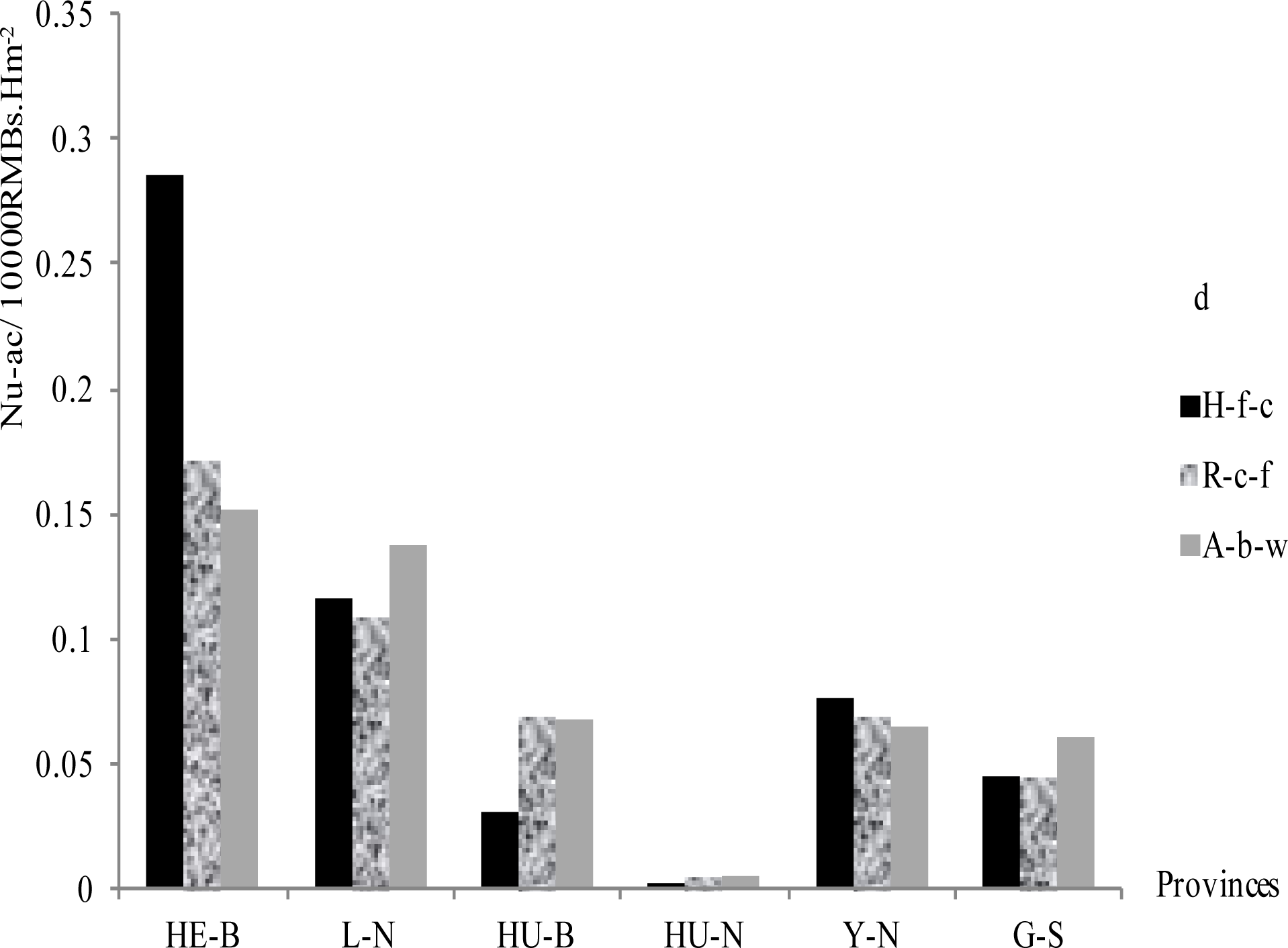
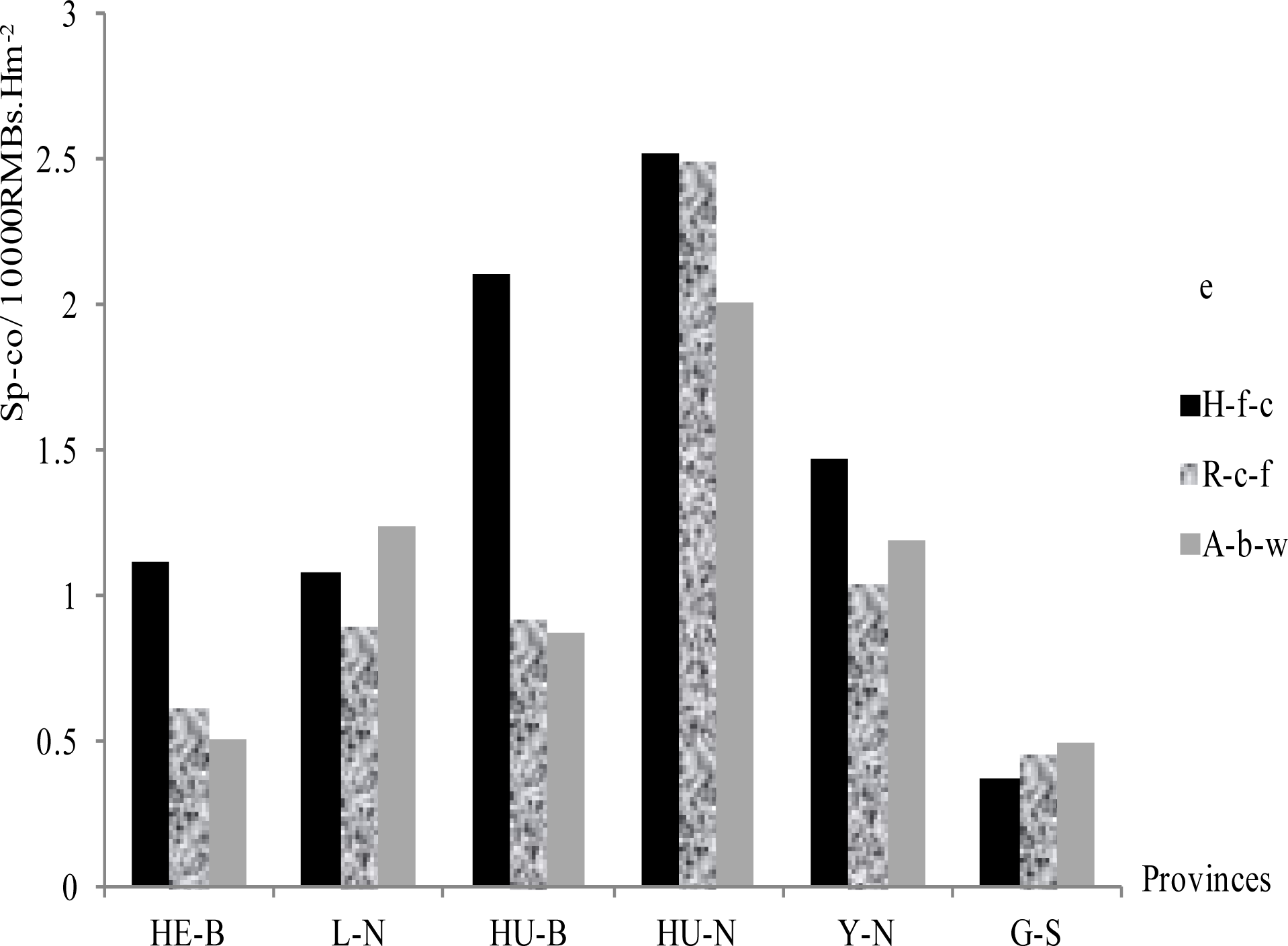
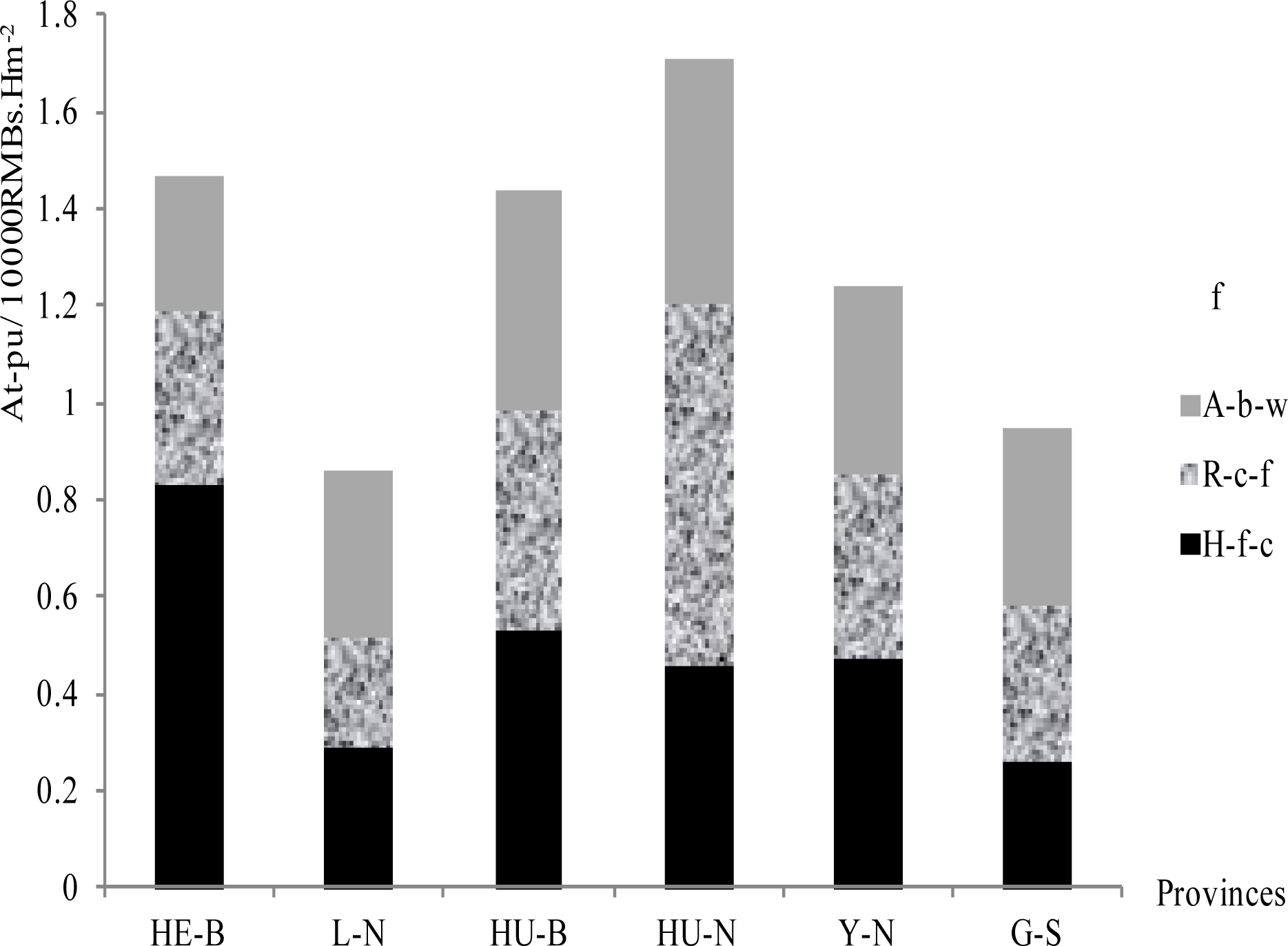
Annual categorized unit area B-Vs of different forest restoration ways.

**Table 4.**
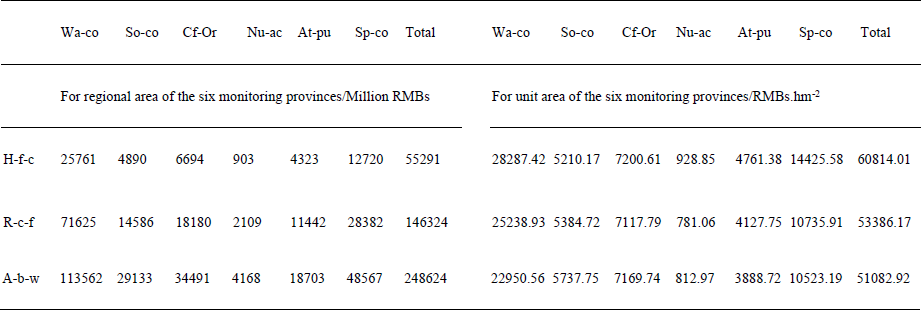
Summary of annual B-Vs in different forest restoration ways.

## 5. Discussions

In this study, we took the CCFP as the object to analyze the features and the correlations of categorized forest ecological benefit values. The results are also of common interest to general forest management. Although there were some studies on the similar topic (Niu et al. 2012; Wang et al. 2011), this study will help to understand more about the details of categorized forest ecological benefit values in the aspects of forest restoration ways, regional specifics. In addition, the results could be of relevance to the environment protection and be used for reference for the future construction of CCFP.

### 5.1. Regional differences of B-Vs

Regional B-Vs are related to the local natural conditions including landform, soil, climate and so on (China’s State Forestry Administration 2014; Sun et al. 2007, 2006; Jackson et al. 2005). We have the consistent results in this study.

The unit area B-Vs of species conservation and water conservation in southern regions (e.g. HU-N, Y-N) were higher than the northern regions (Fig. 3; Fig. 5; Fig. 7). The abundant water and heat resources in southern regions make the forest and other spices easy to make use of the resources for growth and reproduction, and hence the diversity and water-holding ability of these species increased. Only one exception is in HE-B where produced the highest unit area water conservation B-V especially in the ‘hillside forest conservation’ way. The more plateau sandy land and more shrubs in the northern parts in HE-B, and the more raining in its hot season, make the shrubs easy to develop their stronger function on water conservation in the sandy land that suffers from serious water loss.

The larger area of fast-growing forest seems to be the main reason for the highest unit area B-V of nutrient accumulation in HE-B. More hillsides with serious soil erosion make the soil loss more obvious in G-S and Y-N, which might be the reason for having high soil conservation B-Vs, as shown in our study (Fig. 3; Fig. 5; Fig. 7).

According to Yin (2010), the ‘carbon fixation and oxygen release’ B-V of mixed forest is significantly higher than that of coniferous forest (e.g. *Pinus massoniana* Lamb.), and we have the consistent result that HU-N had the lowest ‘carbon fixation and oxygen release’ B-V but produced the highest atmosphere purification B-V with its more coniferous forest. Probably, the low growth rate of coniferous forests resulted in the lowest ‘carbon fixation and oxygen release’ B-V and nutrient accumulation B-V. In addition, the result also indirectly reflected that the coniferous forests might have stronger ability of atmosphere purification. This speculation is contrary to the previous study of Nie et al. (2015), in which they concluded that the purification capacity of different types of urban forest on atmosphere could be ranked as broad-leaved mixed forest > planted bush > conifer forest. Whereas, Shi et al. (2016) presented the same result as we do. They concluded that the ability of atmosphere purification of coniferous forests is higher than broad-leaved forests.

### 5.2. B-V ranks of forest restoration ways

The rank of classified regional B-Vs in the six monitoring provinces was—‘afforestation on suitable barren hills and wasteland’ > ‘returning cropland to forest’ > ‘hillside forest conservation’. Whereas, the rank of average unit area total B-Vs was opposite—‘hillside forest conservation’ > ‘returning cropland to forest’ > ‘afforestation on suitable barren hills and wasteland’ in CCFP (Table 4). It appeared that the regional B-V in the way of ‘afforestation on suitable barren hills and wasteland’ was larger than the other two ways. In fact, this was partly because of its larger area (Fig. 2), and the rank of unit area total B-Vs reflected the reality of the ecosystem service. Shi et al. (2016) concluded that the ecosystem services of ‘returning farmland to forests’ and ‘closing hillsides to facilitate afforestation’ were better than those of ‘afforestation on barren hills and wasteland’. This result gave a further example to illustrate our conclusion. The features of unit area total B-V can be reasonably considered in choosing forest restoration way. Whereas, B-V is not the only goal of CCFP, regional specifics and actual needs should be taken into account to achieve joint-win of ecological, social and economic benefits in the process of forest restoration.

### 5.3. Relationships among the B-Vs

Forests usually cannot simultaneously produce multiple, positive ecosystem services because of the trade-offs among different or competing functions. Maximizing one service may cause substantial declines of other services (Bennett et al. 2009). However, our result showed that forests could produce multiple, consistent positive ecosystem services for the China’s ‘hillside forest conservation’ way on regional scale (Table 1). It is essential to understand the dynamic relationships among all forest ecosystem services (Wang et al. 2011). In addition, Wang et al. (2011) analyzed the relationships between forest cover and runoff on different area scales and concluded that the correlations varied between large scale and meso scale. However, few studies have reported on the correlations among the categorized B-Vs, which were calculated in this study. Our results showed that some of the correlations were consistent and others were not between regional scale and unit area scale. This result could be helpful for considering pros and cons between forest single managing goal and total ecological benefits.

### 5.4. Unit area total B-Vs in CCFP

Wang et al. (2011) released the calculated result of B-V in China general forest ecosystem (40 000 RMBs.hm^−2^.a^−1^–50 000 RMBs.hm^−2^.a^−1^). According to the result of this study, the range of forest annual unit area total B-Vs was 35 000 RMBs.hm^−2^.a^−1^–100 000 RMBs.hm^−2^.a^−1^ in CCFP. The average annual unit area total B-Vs of different forest restoration ways was 51 082 RMBs.hm^−2^.a^−1^–60 814 RMBs.hm^−2^.a^−1^ (Table 4). This result indicated that the unit area total B-Vs in CCFP was larger than that of China general forest ecosystem.

Furthermore, Niu et al. (2012) concluded that the percentages of water values were 40.51% of the total value in Chinese forest ecosystem. Shi et al. (2016) released the percentages of water values were 28.90%, and Qin (2009) released the percentages of water values was 54.09% in CCFP. Our study presented more detailed percentages of water values than the previous in deferent restoration ways. The results were 46.6% on regional scale and 46.5% on unit area scale in ‘hillside forest conservation’ way, 49.0% on regional scale and 47.3% on unit area scale in ‘returning cropland to forest’ way, 45.7% on regional scale and 44.9% on unit area scale in ‘afforestation on barren hills and wasteland’ way in CCFP.

We deduced that CCFP was carried out in the regions with serious ecological degradation, in which their B-Vs were more obvious than the general regions. Up to present, the unified calculation methods of B-V were not formed yet. Different calculation methods might also result in the inconsistencies.

### 5.5. Performances of categorized B-Vs

Water is the most sensitive and limiting ecological factor in forest eco-system (Wang et al. 2011). Our result showed that water conservation B-V was the main part in total B-V in CCFP, and hence, it makes the water more sensitive to the total B-V.

Nutrient accumulation B-V was the least in total B-V and varied in different regions in CCFP. The higher nutrient accumulation B-Vs occurred in the North and Northeast China (Fig. 3, Fig. 5 and Fig. 7). This result might be related to the differences of climate, soil and afforestation species.

Wang et al. (2011) concluded the performance of the overall B-Vs in China’s general forest by the following sequence: water conservation B-V > species conservation B-V > carbon fixation and oxygen release B-V > soil conservation B-V > atmosphere purification B-V > nutrient accumulation B-V. We had the same result in CCFP not only in regional but also in unit area (Table 4).

### 5.6. Limitations

We did not present all the regressions of the B-Vs due to space limitations in the article. There are also many ecological service values for the forest should be taken into the analysis. Such as noise reduction, landscape value and so on. We also did not involve them for lack of the detailed materials of these aspects. Duo to the limitations of the level of cognition and study, some points of the discussion are speculative. We are willing to communicate with our peers to improve our research.

### 6. Conclusions

In the six ecological monitoring provinces in 2013, Water conservation B-V was the highest and nutrient accumulation B-V was the lowest whether on regional or unit area scale in CCFP. The rank of categorized B-Vs was—(water conservation B-V) > (species conservation B-V) > (carbon fixation and oxygen release B-V) > (soil conservation B-V) > (atmosphere purification B-V) > (nutrient accumulation B-V) in CCFP.

In CCFP, forest ecological B-Vs varied in different forest restoration ways and different regions. The rank of average unit area total B-Vs was—‘hillside forest conservation’ > ‘returning cropland to forest’ > ‘afforestation on suitable barren hills and wasteland’. Unit area B-Vs of species conservation and water conservation in southern regions were higher than that of northern and northwestern regions in CCFP. The hot and rainy regions produced higher species conservation B-Vs, and the regions with more coniferous forest had higher atmosphere purification B-Vs and lower ‘carbon fixation and oxygen release’ B-Vs. The regions with more hilly area or more sandy land had higher soil conservation B-Vs. ‘Hillside forest conservation’ was a better way for the regions aiming at water conservation, atmosphere purification and species conservation. For the regions aiming at soil conservation, ‘afforestation on suitable barren hills and wasteland’ was more suitable. The ‘hillside forest conservation’ restoration way and the water conservation B-V should be paid more attention in China’s future forest restoration. We suggest that suitable forest restoration ways should be selective according to the regional specific.

There were correlations among the categorized B-Vs, and the correlations varied with different forest restoration ways in CCFP. Water conservation B-V had significantly positive correlation with the relevant total B-V and positive correlation with the relevant atmosphere purification B-V on both regional scale and unit area scale. Species conservation B-V of unit area was negatively correlated with the relevant nutrient accumulation B-V except in the way of ‘afforestation on suitable barren hills and wasteland’. Regional species conservation B-V had significantly negative correlation with the relevant nutrient accumulation B-V except the ‘hillside forest conservation’ way. Knowing about the correlations among the categorized B-Vs could clarify the targeted restoration ways according to the goal of ecological benefit.

### Abbreviations

CCFP: The China’s Conversion Cropland to Forest Program
CFERN: The Chinese Forest Ecosystem Research Network
B-V: Forest ecological benefit value
Ecological monitoring provinces: 
HE-B: Hebei province
L-N: Liaoning province
HU-B: Hubei province
HU-N: Hunan province
Y-N: Yunnan province
G-S: Gansu province
In Figures and Tables: 
H-f-c: Hillside forest conservation
R-c-f: Returning cropland to forest
A-b-w: Afforestation on suitable barren hills and wasteland
Wa-co: water conservation
Sp-co: Species conservation
At-pu: Atmosphere purification
So-co: Soil conservation
Cf-Or: Carbon fixation and oxygen release
Nu-ac: Nutrient accumulation

## Declarations

### Acknowledgements

The Hebei Provincial Science & Technology Supporting Program (No.15227652D) and CFERN & BEJING TECHNO SOLUSIONS

Award Funds on excellent academic achievements provided the Project support. The work was also guided by ‘Observation Methodology for Long-term Forest Ecosystem Research’ of National Standards of the People’s Republic of China (*GB/T 33027–2016*).

We appreciate that Dr. Bing Wang, a researcher from Chinese Academy of Forestry provided some of the technical data. The author is also indebted to Dr. Yuwu Li from Xishuangbanna Tropical Botanical Garden, Chinese Academy of Sciences for his advice on the article. In addition, Key Lab. of Genetic Resources of Forest and Forest Protection of Hebei Province should be credited for part of its Supporting. We thank the journal reviewers for their detailed and the constructive comments on the manuscript.

### Availability of data and materials

We declare that the materials described in the manuscript, including all relevant raw data, are freely available, without breaching participant confidentiality.

### Legal statement

All research work reported in this study was performed in accordance with all relevant legislation and guidelines.

### Authors’ contributions

All authors conceived the study performed research and analyzed data. Wen-Ge Yuan wrote the paper.

### Ethics approval and consent to participate

This manuscript does not report on or involve the use of any animal or human data or tissue.

### Consent for publication

This manuscript does not involve animal or human study. We understand that the text and any pictures published in the article will be freely available on the internet and may be seen by the public.

### Authors’ information

Wen-Ge Yuan ^a, b, c^ Jian-Wei Zheng _a, c_ Jian-Cai Gu _a, c_* Gui-Qiao Lu ^a, c^

^a^ Forestry College, Agriculture University of Hebei, No. 2596, Southern street of Lekai, Baoding 071000, China

^b^ Langfang Academy of Agriculture and Forestry Sciences, No. 285, Guangyang Road, Langfang 065000, China

^c^ Key Lab. of Genetic Resources of Forest and Forest Protection of Hebei Province, Baoding 071000, China

